# Spring mineral water-borne bacteria reshape gut microbiota profiles and confer health benefits

**DOI:** 10.1101/433821

**Authors:** YP Chen, LL Tan, DM Chen, Q Xu, JP Song, QP Zeng

## Abstract

**Background:** Although dietary patterns are recognized to affect health by interfering with gut microbiota homeostasis, whether live or dead bacteria-bearing spring mineral water (MW) would also exert beneficial effects on health upon curing gut dysbiosis remains unknown.

**Results:** Due to harboring live bacteria, the heated but unboiled MW from Bama, where centenarians are ubiquitously inhabited, reshapes the gut microbiota from a traveler-type to a local resident-type except for *Prevotella*. While chondroitin sulfate, a component occurring in livestock and poultry meats, increases the richness of sulfatase-secreting bacteria and sulfate-reducing bacteria, Bama MW dampens the overgrowth of those colon-thinning bacteria and hampers the overexpression of multiple genes responsible for anti-inflammation, anti-oxidation, anti-hypoxia, anti-mutagenesis, and anti-tumorigenesis.

**Conclusions:** Bama spring MW prevents the early-phase onset of breast cancer by curating gut dysbiosis. MW also compromises chromosomal DNA damage and ameliorate mitochondrial dysfunctions, implying it may extend lifespan.

## Background

Obesity and overweight are the most pivotal factors predisposing the development of type 2 diabetes (T2D) and breast cancer. In 2015, 600 million obese adults and 100 million obese children were announced from 195 countries [1]. As of 2015 there were 392 million people diagnosed T2D worldwide [2]. In 2012, 1.68 million new cases of breast cancer and 522,000 deaths subjected from breast cancer were recorded by the World Health Organization (WHO) [3].

It was also known that red meat-contained chondroitin sulfate (CS) can flourish the sulfatase-secreting bacterium (SSB) *Bacteroides thetaiotaomicron* and the sulfate-reducing bacterium (SRB) *Desulfovibrio piger* [4]. For a diet depleting fibers, the mucin-eroding bacterium (MEB) *Akkermansia muciniphila* becomes abundant and starts to degrade the mucin that covers the surface of colonic linings [5], by which SSB and MEB release sulfate, and SRB reduce sulfate to hydrogen sulfide.

Although *Akkermansia muciniphila* were considered beneficial to mucin turnover [6], its overgrowth may also compromise the colonic integrity and enhance the enteric permeability. As results, gut microbes or their components such as lipopolysaccharide (LPS) can simply trespass the colonic barrier and enter the blood circulation to trigger immune activation and inflammatory responses. Therefore, *Akkermansia muciniphila* would be harmful if bacterial degradation of mucin-derived nutrients exceeds production [4, 5].

Given that the abundant SSB and MEB, through opening the mucus barrier and leaking the bacterial endotoxin, serve as the etiological initiators of T2D [7], it seems that an alteration of the dietary patterns from a high-fat diet to a high-fiber diet should improve this unhealthy condition. A case-cohort study conducted in 8 European countries indicated that dietary fiber intake is associated with a lower risk of T2D [8]. Most recently, dietary fibers that enrich the bacteria producing short-chain fatty acid (SCFA) were attributed to the alleviation of T2D [9].

In similar, dietary factors have been linked to the increased risk of breast cancer, including a high-fat diet [10] and an excessive alcohol intake [11]. Although the meta-analysis trying to link fiber intake with breast cancer gave rise to the mixed results [12], a tentative association of low fiber intake during adolescence with low breast cancer incidence was established [13]. Besides, the dietary heme rich in red meat was demonstrated to induce epithelial hyper proliferation via enhancing gut microbiota disturbance (dysbiosis) [14].

Alternatively, mineral water (MW) was addressed to affect human health in addition to diets. For instance, MW rich in sodium was found to increase the insulin sensitivity in postmenopausal women [15]. Consumption of the bicarbonate-rich MW was shown to prevent T2D by increasing the lean-inducible bacteria *Christensenellaceae*. Previous studies were actually documented that *Christensenellaceae* can be enriched in lean group (BMI<25) as compared with obese group (BMI>30) [16]. It was also reported that transplantation of *Christensenella minutia* to germ-free mice can reduce weight gain. Moreover, the abundance of *Dehalobacteriaceae* that are increased after MW drinking were found to positively correlate the abundance of *Christensenellaceae* [17].

However, it remains unknown whether the natural and fresh MW would be implicated in human health such as lifespan extension and breast cancer prevention via waterborne bacteria *per se*. Herein, we report for the first time that the unboiled or bottled MW from Bama, a well-known longevity county located in South China, can reshape the gut microbiota communities in both murine and human. Consequently, Bama MW drinking blocks mouse mammary hyperplasia accompanying with downregulation of the breast cancer cell proliferation marker Ki67 [18] and the triple-negative breast cancer (TNBC)-specific transcription factor BCL11A [19].

Through anti-oxidation, anti-mutagenesis and anti-tumorigenesis, Bama MW drinking may compromise chromosomal DNA damage and ameliorate mitochondrial dysfunctions, thereby reducing morbidity and mortality from metabolic diseases. Our results may partially explain why centenarians, who have a life-long habit of fresh MW drinking, were frequently seen in Bama. Our new findings are most likely provide a non-medical option for hampering the metabolic diseases alternative to high-fiber diet consumption.

## Methods

### Mouse modeling and treatment

The SPF female BALB/c mice (18-22 g) were provided by The Experimental Animal Centre of Guangzhou University of Chinese Medicine in China (Certificate No. 44005800001448). Animals were housed on a 12-h light and 12-h dark cycle at 25°C with *ad libitum* (AL) chow and free tap water drinking. After 3-week quarantine, animals were randomly divided into 6 groups: (1) a group of AL mice; (2) a group of mice intragastrically administered 1g/kg CS (FocusChem, Shandong, China) every other day for 6 months; (3) a group of mice intragastrically administered 1g/kg CS every other day and 10^8^ *Bacillus cereus* (BC, Huankai Microbial Sci & Tech Co Ltd., Guangdong, China) 4 times during 6-month modeling (CS+BC mice); (4) a group of mice peritoneally injected 0.25mg/kg LPS (Sigma Aldrich) every other day within 3 months (LPS mice); (5) a group of CS-BC mice with the bottled Bama MW drinking (CS-BC+MW) started from modeling (for 6-month drinking); (6) a group of CS-BC mice with the boiled MW drinking (CS-BC+BMW) started from modeling (for 6-month drinking). The bottled Bama MW with an omitted trademark was purchased from the market, and the boiled Bama MW was prepared from the bottled Bama MW. Mouse fecal samples were collected from the mouse cage (1 g for one test).

### Diet consumption, MW drinking and fecal sampling from volunteers

Two travelers of one male aged 57 and one female aged 20 were selected, and 8 local residents of 4 males and 4 females aged from 10 to 80 years old were enrolled (with anonymous and informed consent). The volunteers were allowed to the unrestricted diet consumption, among which the travelers drank the heated but unboiled (at <60°C for 1 minute, resembling the pasteurization of yogurt) MW daily for 4 weeks, and the local residents drank the unprocessed MW drained from the mountain, and the natural MW drunk by the travelers was fetched from a mouth of the spring located in Baimo Village of Bama County, Guangxi, China.

Using sampling tubes provided by TinyGene, Shanghai, China, fecal samples were collected from local residents and travelers. The former samples were only collected for once, and the latter samples were collected for once prior to MW drinking and for 4 times post MW drinking. The *in vitro* aerobic culture of bacteria in the unboiled and bottled MW was performed by TinyGene, Shanghai, China obeying a standard procedure.

### Gut microbiota metagenomic analysis

The gut microbiota in fecal samples or the unboiled and bottled MW was classified by the high-throughput 16S VX region sequencing. DNA extraction and detection, amplicon purification, library construction, online sequencing, and data analysis including paired end-reads assembly and quality control, operational taxonomic units clustering and species annotation, alpha and beta diversity of mouse fecal samples were conducted by Novogene, Beijing, China; and those similar tests of human fecal samples were conducted by TinyGene, Shanghai, China.

### Enzyme-linked immunosorbent assay (ELISA)

ELISA kits for mouse TNF-α, IL-1β, VEGF, Mn-SOD, BRCA1, and bacterial LPS were purchased from Andy Gene, Beijing, China. All tests were conducted according to the manufacture’s manuals.

### Quantitative polymerase chain reaction (qPCR)

RNA isolation, purification, electrophoresis, reverse transcription, and quantification were performed obeying the standard protocol. The relative copy numbers = 2^−ΔΔCт^, in which ΔCт = Ct target gene − Ct reference gene, ΔΔCт = ΔC treatment sample − ΔC control sample. The primers listed in Table 1 for mice were designed and applied under the following amplification condition: 95°C, 60s; and 95°C, 30s, 60°C, 35s for 40 cycles.

**Table 1.**
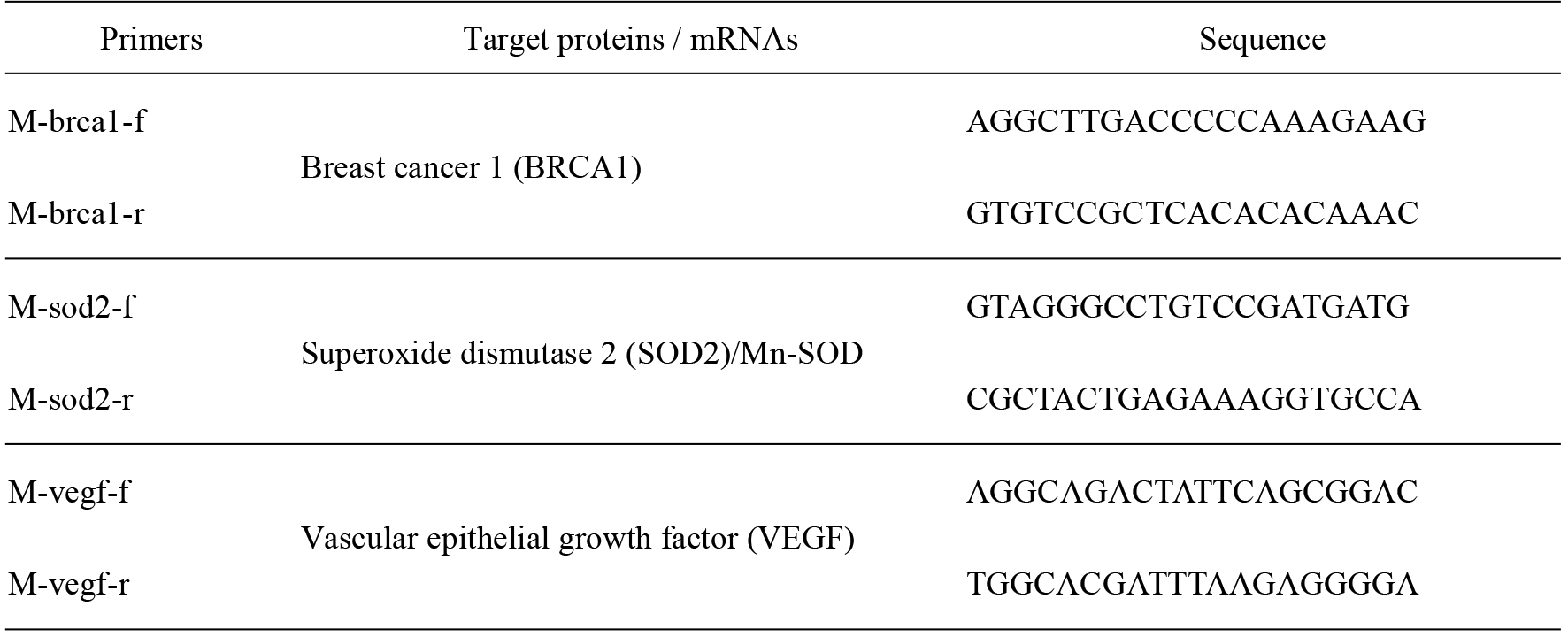
Primer sequences for amplification of mouse target mRNAs by RT-PCR

### Histochemical analysis

Fix the dissected tissue by immersion into a 10% formalin solution for 4 to 8 hours at room temperature. Mount in OCT embedding compound, and freeze at −20 to −80°C. Cut 5-15 μm thick tissue sections using a cryostat. Thaw-mount the sections onto gelatin-coated histological slides. Slides are pre-coated with gelatin to enhance adhesion of the tissue. Dry the slides for 30 min on a slide warmer at 37°C. Sections were deparaffinized by xylene, re-hydrated by gradient alcohol, and washed in distilled water. After haematoxylin staining, wash in running tap water, differentiate in 1% acid alcohol, wash again in running tap water, blue in 1% ammonia, wash again in running tap water, and rinse in 95% alcohol. After eosin counter staining, dehydrate through gradient alcohol, clear in xylene, and mount with xylene-based mounting medium.

### Statistical analysis

The software SPSS 22.0 was employed to analyze data, and the software GraphPad Prism 6.01 was employed to plot graphs. The Independent Simple Test was used to compare all groups, but the Kruskal-Wallis Test followed by Nemenyi test was used when the data distribution is skewed. The significance level (p value) was set at <0.05 (*), <0.01 (**), <0.001 (***) and <0.0001 (****).

## Results

### Gut bacterial reshaping from traveler-individual to local resident-special by unboiled MW drinking

To disclose the secret of health and longevity in Bama citizens, who have a life-long habit of fresh MW drinking, we classified the representative gut bacterial profiles of the randomly enrolled local residents, with the males and females aged from 10 to 80 years old. Interestingly, their gut bacteria show the marked sexual difference on the phylum level. The males carry 4 major phyla, accounting for 2/3 of *Bacteroidetes*, and 1/3 of *Firmicutes*, *Proteobacteria* and *Fusobacteria*, whereas the females possess 3 major phyla, including 2/3-1/2 of *Bacteroidetes*, and 1/3-1/2 of *Firmicutes* and *Proteobacteria* (Fig.1a). It was unknown why *Fusobacteria* are only present in the males, but absent in the females.

**Figure 1.**
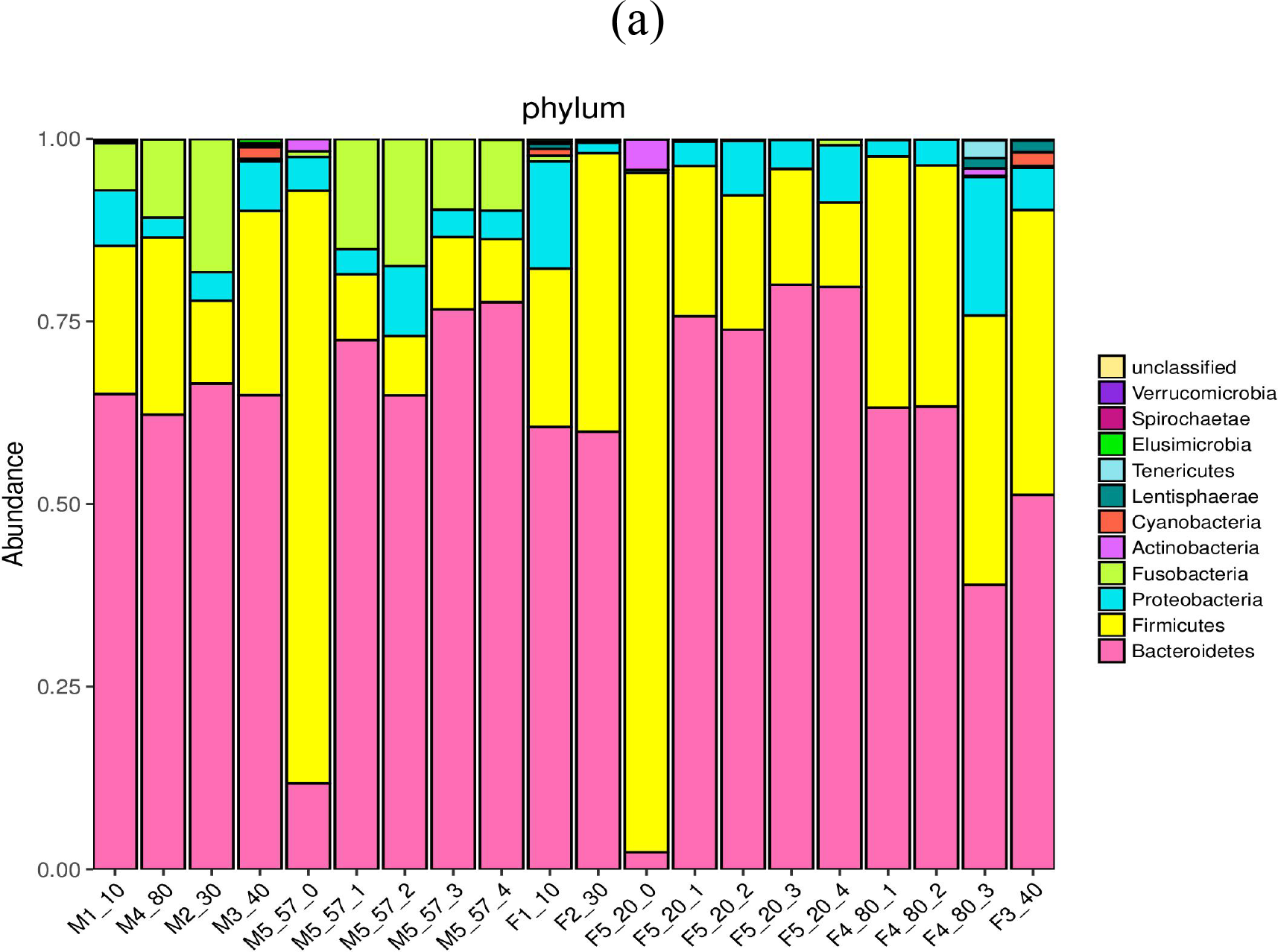
The phylogeny heat-map of gut bacterial phyla (a) and genera (b) and PCA (c) in the local residents and travelers. M represents male; F represents female; the numbers following M/F represent the representative groups, age ranges, and sampling frequencies or individual sampling, respectively.

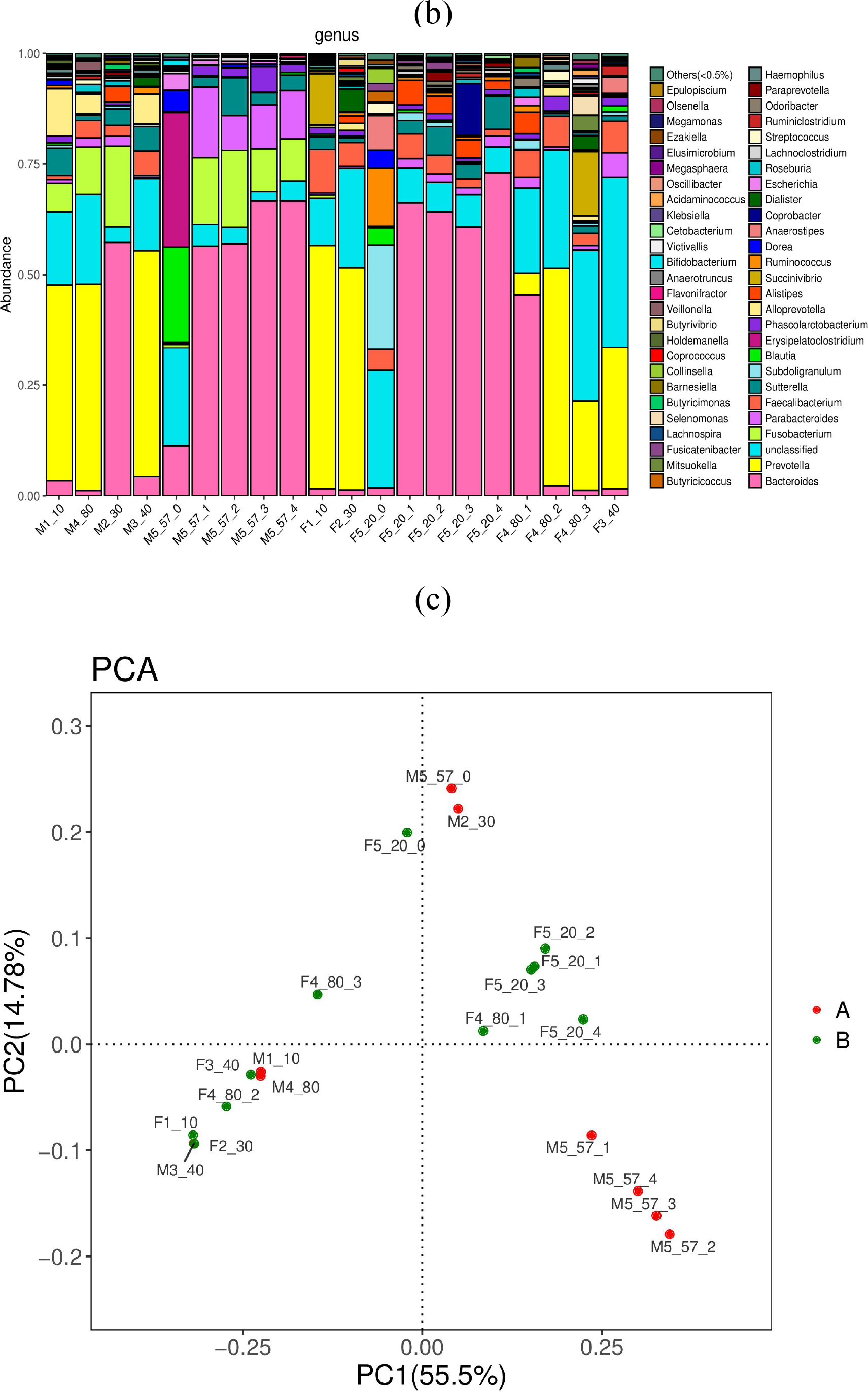

Simultaneously, the dynamic changes of gut bacterial compositions prior to and post MW drinking were monitored in two travelers aged 57 and 20 years old, respectively. As results, the gut bacterial phyla of travelers are completely different from those of local residents. For example, the travelers’ gut bacteria comprise of the most abundant *Firmicutes*, whereas the local residents’ gut bacteria are predominated by the most abundant *Bacteroidetes*. After drinking the heated but unboiled MW, however, the gut bacteria of travelers are quite similar with that of local residents (Fig.1a).

By comparison among the bacterial phyla, it could be concluded that the unboiled MW may extraordinarily decrease the abundance of *Firmicutes*, which predominantly exist in the gut of travelers, as well as extremely increase the richness of *Bacteroidetes*, which are overwhelming in the gut of local residents. Additionally, many unclassified genera are present in the local residents’ gut, but absent in the travelers’gut (Fig. 1b).

From the principle component analysis (PCA) (Fig. 1c), it was noted that the effectors on the gut bacterial composition are similar in the local residents, so their operational taxonomic units (OTUs) are distributed together at the left-bottom quadrant. Interestingly, OTUs from the female traveler sample prior to MW drinking (F5_20_0) is distributed at the different left-top quadrant without identity to the local residents (except for the M2_30 who might be an immigrant), whereas OTUs from her sample after MW drinking (F5_20_1, _2, _3 and _4) become very closed to a local resident sample (M4_80_1).

From comparison of the bacterial genera, it was noticed that the local residents’ gut-specific *Prevotella* are absent in the travelers’ gut even after daily MW drinking for 4 weeks. Although local residents and travelers commonly possess *Bacteroidetes*, the local resident-specific *Prevotella* cannot replace *Bacteroides* through MW drinking, suggesting that *Prevotella* should be enriched by other factors such as the diets.

It was also significant that the gut bacterial composition of one sample from the boy (M1_10) and one sample from the girl (F1_10) show an extraordinarily higher proportion of *Prevotella* and an extremely lower proportion of *Bacteroides*. Interestingly, one sample from the 80-years-old man (M4_80) and two samples from the 80-years-old women (F4_80_2 and F4_80_3) also exhibit a high ratio of *Prevotella* to *Bacteroides*, which is similar with the gut bacterial floras of children, and may represent the Bama-specific gut microbiome profiles (Table 1).

**Table 1.**
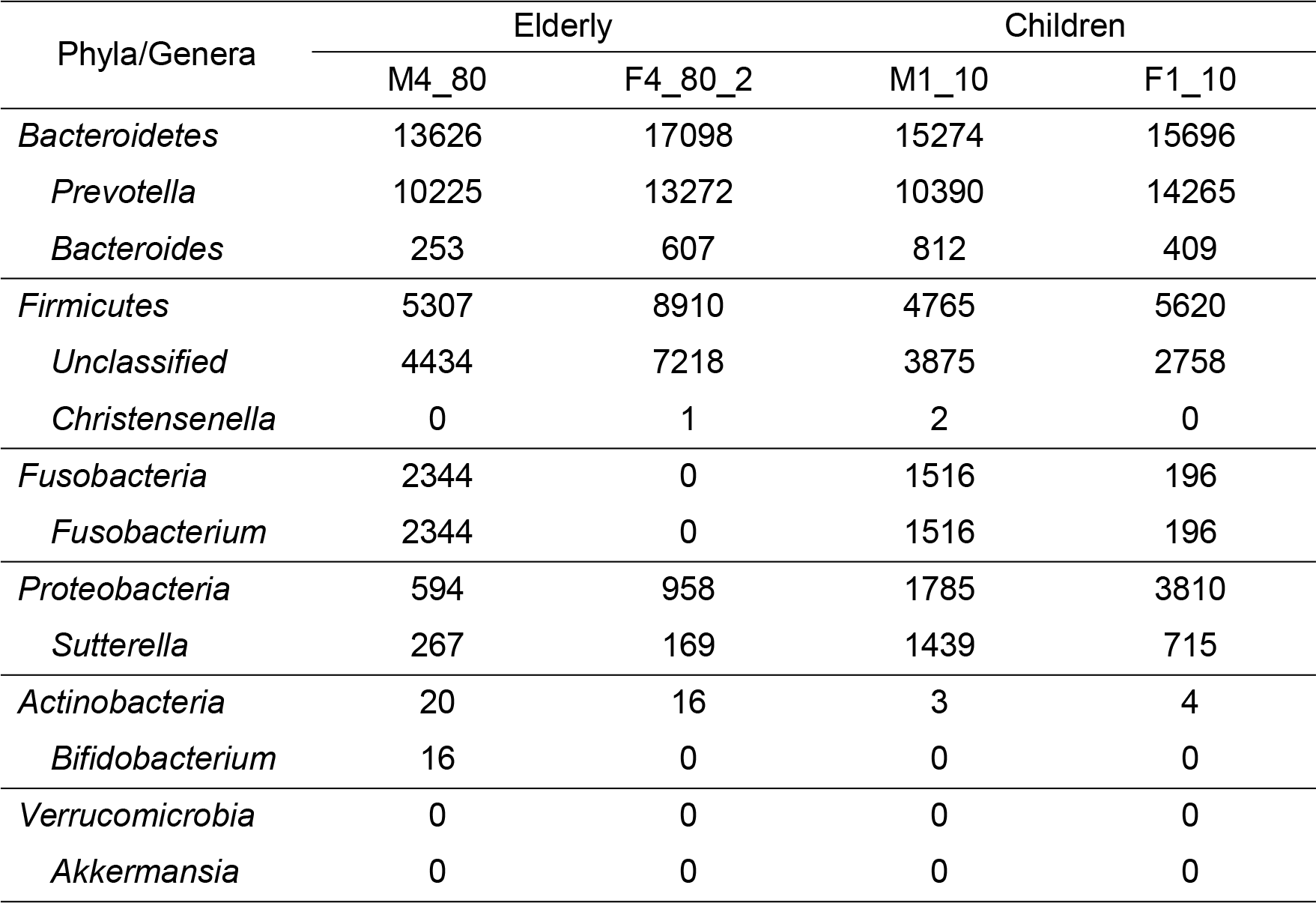
Comparison of the sequence counts in the selective gut bacterial genera between Bama elderly and children

Additionally, samples from either elderly or children have the equal sequence counts of an unclassified genus belonging to the class *Clostridia* in *Firmicutes* as well as the genus *Sutterella* in *Proteobacterium*. As to the genus *Fusobacterium* in *Fusobacteria*, the female samples have less bacterial sequence counts than the male samples with an unknown reason. Surprisingly, the so-called lean-related bacterial genus *Christensenella*, almost absent in the gut of local residents.

It was also noted that the sequence counts from the well-known prebiotics, *Bifidobacterium* and *Akkermansia*, are from few in elderly to zero in children, implying only null or undetectable levels of these MEB might be beneficial by facilitating mucin turnover, otherwise an abundant presence of MEB would be harmful due to mucin depletion.

### *In vitro* culture and metagenomic sequencing of bacteria in the natural MW samples

To understand the mechanism underlying how can MW reshape the gut bacterial community, either by inhibiting bacterial growth or replacing the original gut bacteria, we classified the bacterial taxa in the natural MW samples with or without *in vitro* culture. A genera heat-map of bacteria available from the uncultured and cultured MW samples was illustrated in Fig. 2a.

**Figure 2.**
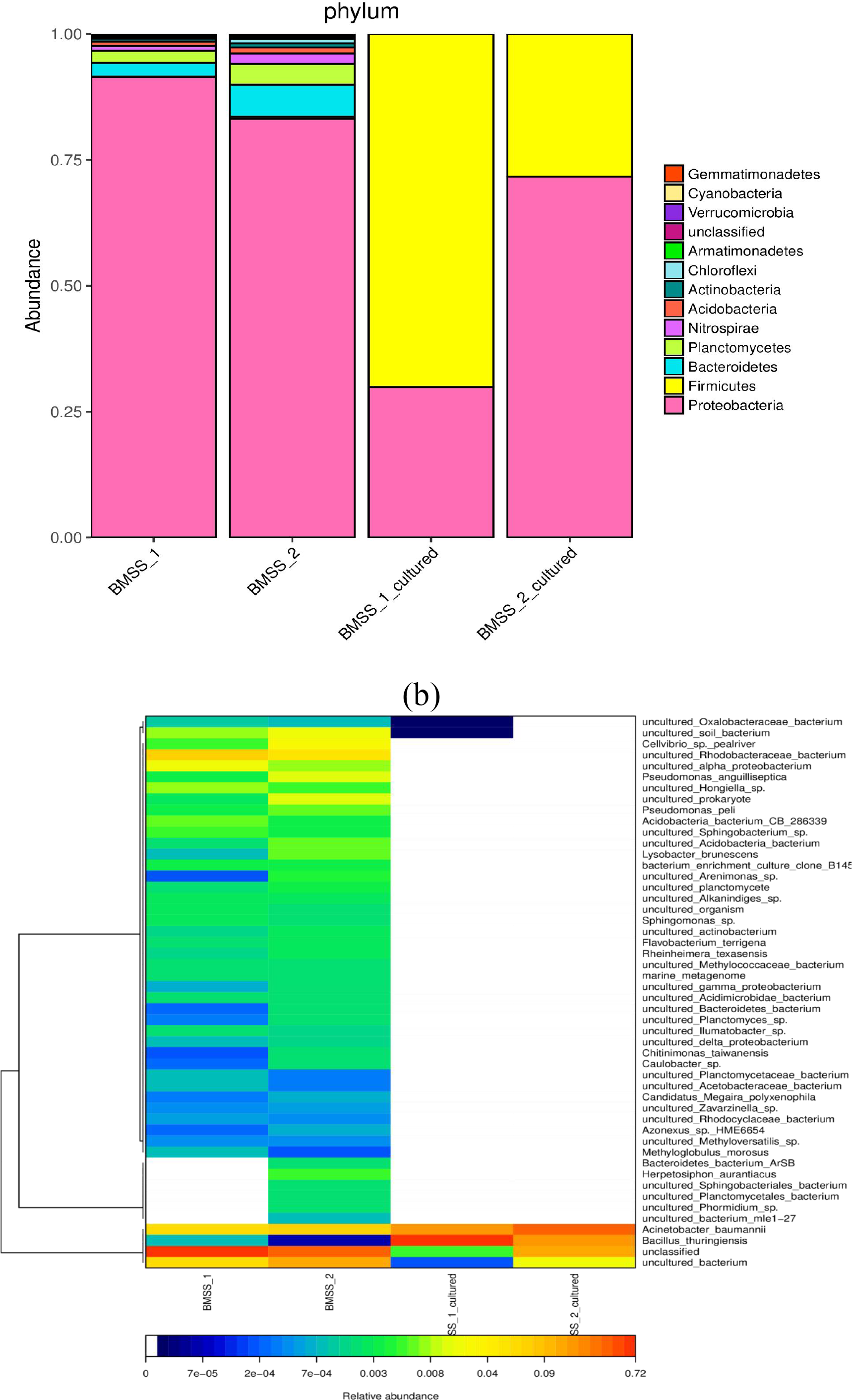
The taxonomic heat-map of bacterial phyla (a) and the ranked heat-map of bacterial species (b) in the uncultured and cultured MW. BMSS_1 and BMSS_2 represent the bacterial profiles in two uncultured MW samples; BMSS_1_cultured and BMSS_2_cultured represent the bacterial profiles in two cultured MW samples.

The uncultured MW samples contain the similar bacterial phyla to the human gut, including *Proteobacteria*, *Bacteroidetes*, and *Firmicutes* albeit in a distinct proportion. They are characterized by many soil, water, and plant-specific genera of bacteria, such as *Planctomycetes*, *Nitrospirae*, and *Acidobacteria*. The aerobic culture of MW samples, however, gave rise to only two phyla of bacteria, *Proteobacteria* and *Firmicutes*, indicating that a large part of bacteria was uncultivable under the presently chosen aerobic cultural condition (Fig. 2a).

From the phylogeny tree of bacterial species illustrated in Fig. 2b, it was clear that the waterborne bacterial species are less similar with the gut-residing bacterial species although *alpha*-, *gamma*-, and *delta*- *proteobacterium* were found in the natural MW. The most abundant bacterial species emerging in both uncultured and cultured MW samples is *Acinetobacterium baumannii*.

It was evident that the natural MW-borne bacteria might inhibit gut bacterial growth rather than replace gut bacteria because MW-borne bacteria are different from gut-dwelling bacteria. Importantly, no human pathogenic bacteria were identified from the natural MW samples although *Pseudomonas anguilliseptica* is pathogenic to fish (Fig. 2b).

### *In vitro* culture and metagenomic sequencing of bacteria in the bottled MW samples

We also performed *in vitro* culture of the bottled MW sample under the aerobic condition. Unfortunately, no colonies were propagated in cultural plates, indicating bacteria in the bottled MW being mostly dead. However, it could not completely exclude that some sporous bacteria are still survived from the tailored detoxifying procedure, such as ozone exposure and ultraviolet irradiation. On the other hand, uncultivable under the aerobic condition was by no means said to germ-free because many species of bacteria are anaerobically cultivable.

Nevertheless, bacterial DNA, either intracellular or intercellular, suspended in the bottled MW could be sequenced. Notably, a huge number of bacteria (68325 counts), mainly including *Proteobacteria* (51259 counts), *Firmicutes* (9401 counts), and *Bacteroidetes* (6671 counts), were detected in the bottled MW sample. Fig. 3a illustrated the bacterial phyla identified from the bottled MW sample. The most abundant phylum is *Proteobacteria* (0.75), and the relative abundance of other two phyla, *Bacteroidetes* and *Firmicutes* are less than 0.25.

**Figure 3.**
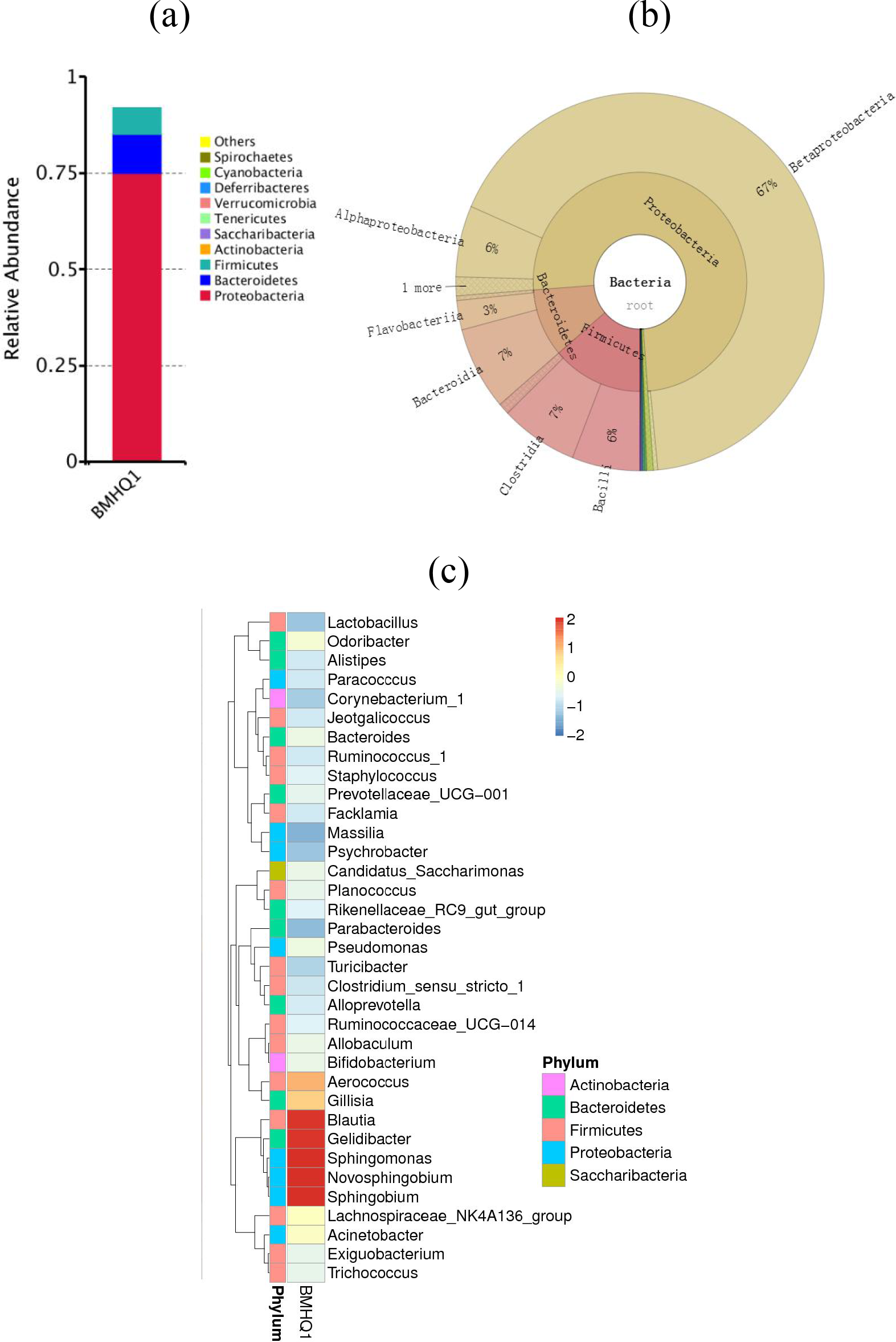
The bacterial phylum composition (a), class proportion (b) and the ranked Top 35 genus heatmap (c) in the bottled MW sample. BMHQ1: a bottled MW sample.

In the level of bacterial classes, *Proteobacteria* mainly include the class *Betaproteobacteria* (0.67) and *Alphaproteobacteria* (0.06). For other major classes, *Bacteroidetes* contain *Bacteroidia* and *Flavobacteria*, and *Firmicutes* contain *Bacilli* and *Clostridia* (Fig. 3b).

From Fig. 3c, it was noticed that the most abundant *Proteobacteria* are the genera *Sphingomonas*, *Novosphingobium* and *Sphingobium. Gelidibacter* and *Blautia* are the abundant genera in *Bacteroidetes* and *Firmicutes*, respectively.

From the proportions of *Proteobacteria* (0.75), *Firmicutes* (0.14) and *Bacteroidetes* (0.10) in the bottled MW samples, compared to that of *Proteobacteria* (0.91), *Firmicutes* (0.0008) and *Bacteroidetes* (0.027) in the natural MW sample, it was thought that the bottled MW might be contaminated though detoxified thereafter by the industrial filtration or irradiation processing.

### Gut fungal compositions in the pre- and post-treated mice

To reveal the gut fungal compositions in mice before modeling, we analyzed the composition of gut fungi in the fecal samples from AL mice. Each mouse has its unique fungal phyla. For example, AL1 mouse has the abundant phyla, *Basidiomycota* and *Zygomycota*, while AL3 mouse shows another abundant phylum, *Ascomycota* (Fig. 4).

**Figure 4.**
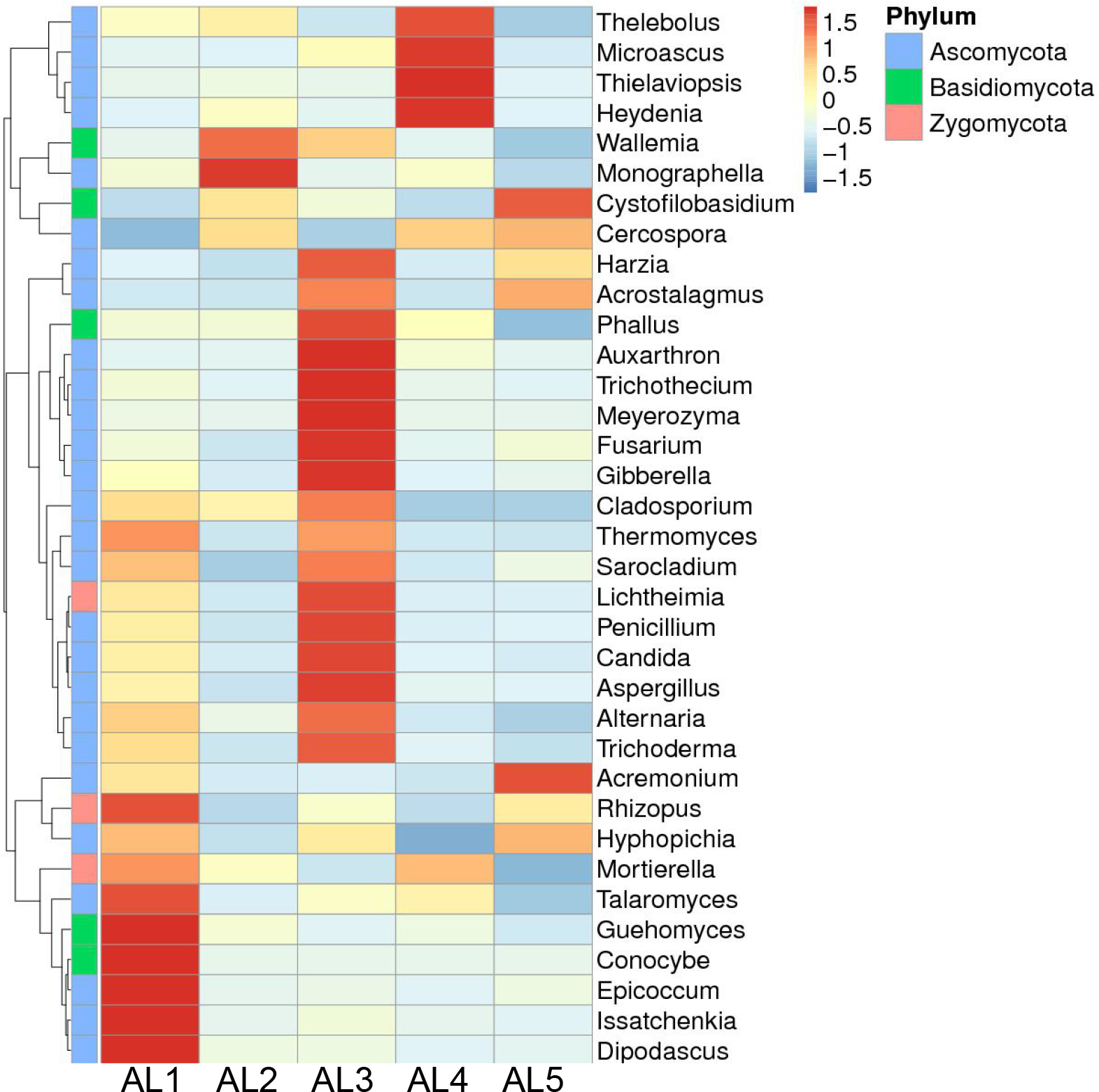
The ranked Top 35 gut fungal genus heatmap of 5 fecal samples from AL mice. AL1-5 represent 5 fecal samples from AL mice.

However, we did not extract and amplify any fungal DNA sequences from the bottled Bama MW. Importantly, we also could not detect fungal DNA in the fecal samples from CS-BC, and CS-BC+MW mice, suggesting that the modeling and drinking procedures impede the survival of fungi in the anaerobic gut niche in mice.

### Comparison of gut bacterial compositions among mice with co-modeling MW drinking

The original intention was to choose the synchronous CS and BC feeding for establishment of the gut dysbiosis model in mice. Because CS can be degraded by BC, we anticipated that *Firmicutes* should be more abundant than other major phyla, *Bacteroidetes* and *Proteobacteria*, after CS-BC feeding. As prediction, *Firmicutes* are enriched, but no significant difference was observed between AL and CS-BC mice (Table 3).

**Table 3.** The proportion of *Firmicutes*, *Bacteroidetes*, and *Proteobacteria* in fecal samples from AL and CS-BC mice.

| Group | *Firmicutes* |  | *Bacteroidetes* |  | *Proteobacteria* |  |
| --- | --- | --- | --- | --- | --- | --- |
|  | AL | CS-BC | AL | CS-BC | AL | CS-BC |
| 1 | 0.48 | 0.56 | 0.42 | 0.37 | 0.08 | 0.04 |
| 2 | 0.47 | 0.80 | 0.43 | 0.12 | 0.07 | 0.06 |
| 3 | 0.42 | 0.50 | 0.49 | 0.42 | 0.06 | 0.05 |
| 4 | 0.50 | 0.55 | 0.45 | 0.37 | 0.04 | 0.04 |
| 5 | 0.40 | 0.51 | 0.53 | 0.42 | 0.05 | 0.04 |
| $\bar{x} \pm s$ | 0.45±0.04 | 0.58±0.11 | 0.46±0.04 | 0.34±0.11 | 0.06±0.01 | 0.05±0.01 |

Note: AL: *Ad libitum* chew-fed mice; CS-BC: chondroitin sulfate and *Bacillus cereus*-fed mice.

To further verify whether CS-BC feeding would increase the richness of the class *Bacilli* in *Firmicutes*, we compared the proportion of *Bacilli* between AL and CS-BC fecal samples. Surprisingly, *Bacilli* even become less abundant in CS-BC samples compared to AL samples. To find out the possible reason why BC feeding does not increase the abundance of *Bacilli*, we compared the proportion of another class, *Clostridia* in *Firmicutes*. Intriguingly, it was noticed that *Bacilli* are even abundant in AL mice, but become scarce in CS-BC mice. In contrast, *Clostridia* are scarce in AL mice, but become abundant in CS-BC mice, as shown in Table 4. These results implied that BC might serve as a competitor to undermine the growth of its counterpart members in *Bacilli*, which could in turn prompt the propagation of *Clostridia*.

**Table 4.**
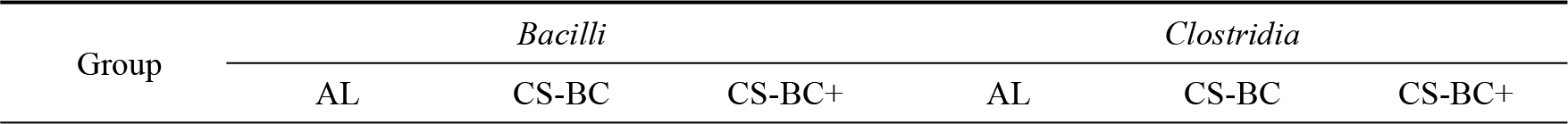
Comparison of the proportion of *Bacilli* and *Clostridia* in the fecal samples from AL, CS-BC, CS+BC+MW/BMW mice

|  | MW/BMW |  |  |  | MW/BMW |  |
| --- | --- | --- | --- | --- | --- | --- |
| - MW/-BMW | 0.39 | 0.09 | 0.11 | 0.06 | 0.66 | 0.12 |
| +MW | 0.31 | 0.09 | 0.21 | 0.11 | 0.39 | 0.04 |
| +BMW | 0.19 | 0.07 | 0.07 | 0.13 | 0.34 | 0.58 |
| $\bar{x} \pm s$ | 0.30±0.10 | 0.08±0.01* | 0.13±0.07* | 0.10±0.04 | 0.46±0.17* | 0.25±0.29 |

Note: AL: *Ad libitum* chew-fed mice; CS-BC: chondroitin sulfate and *Bacillus cereus*-fed mice; CS-BC+MW/BMW: CS-BC mice treated by MW or BMW.

In CS-BC+MW mice, it was observed that *Bacilli* are normalized to abundance again, and *Clostridia* are normalized to less abundance. In contrast, neither increase of *Bacilli* nor decrease of *Clostridia* was seen in CS-BC+BMW mice. This result suggested that the curation of gut dysbiosis by MW drinking may depend upon the presence of some unknown functional competitors, perhaps the heat-sensitive and structure-fragile molecules.

To confirm the assumption that BC can compete with other related bacteria, we compared the richness changes within the phylum *Firmicutes*, the class *Bacilli*, and the genus *Bacillus* in AL and CS mice. Without BC feeding, CS feeding alone increases the abundance of *Firmicutes*, *Bacilli*, and *Bacillus* in CS mice compared to AL mice (Table 5), suggesting CS increases but BC decreases the abundance of *Bacillus*.

**Table 5.**
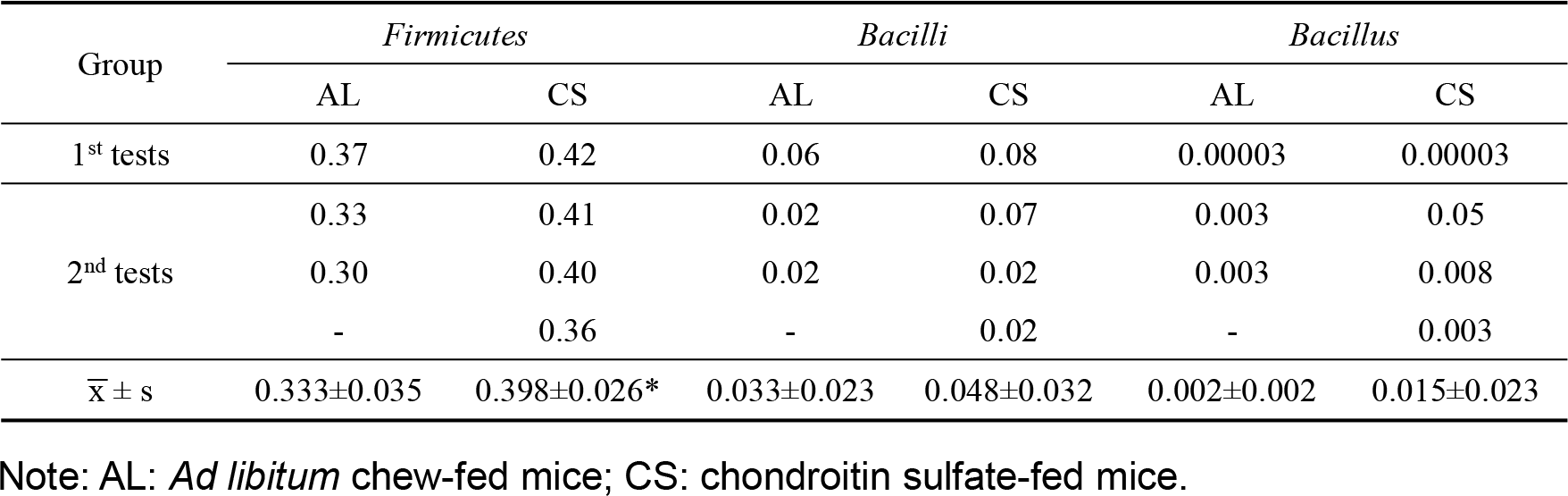
Comparison of the percentages of *Firmicutes*, *Bacilli* and *Bacillus* in the fecal samples from AL and CS mice

### Dynamic changes of mouse gut SSB/MEB and SRB abundance upon co-modeling MW drinking

Because CS is a kind of sulfate that can be degraded by SSB/MEB, it could be anticipated that CS feeding should flourish SSB/MEB and SRB. It was worthy noting that although *Bacteroides* was included in the dynamic analysis of bacterial abundance, not all species in *Bacteroides* are SSB/MEB. For example, *B. thetaiotaomicron* belongs to SSB, and *B. caccae* belongs to MEB.

When compared the bacterial genera of mice pre-modeling, it was only found that *Bacteroides* (containing SSB/MEB) are enriched in one mouse gut and *Bifidobacterium* (MEB) are enriched in another mouse gut (Fig. 5a). After modeling, the bacterial genera belonging to SSB/MEB and SRB are increased to *Rikenella*, *Akkermansia* and *Desulfovibrio* in addition to *Bacteroides* and *Bifidobacterium* (Fig. 5b). In CS-BC+MW mice, the bacterial genera belonging to SSB/MEB and SRB become less abundant. In contrary, the abundance of SSB/MEB and SRB is dramatically increased in CS-BC+BMW mice (Fig. 5c).

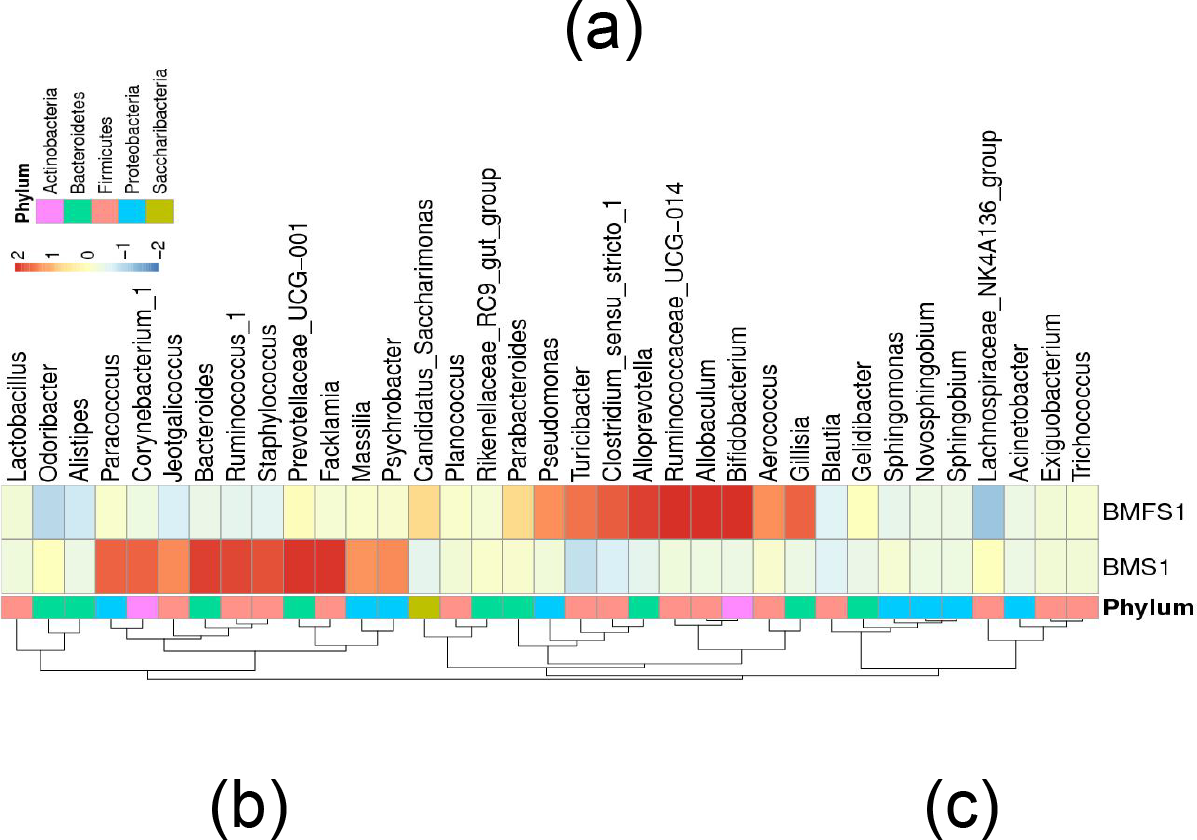

**Figure 5.**
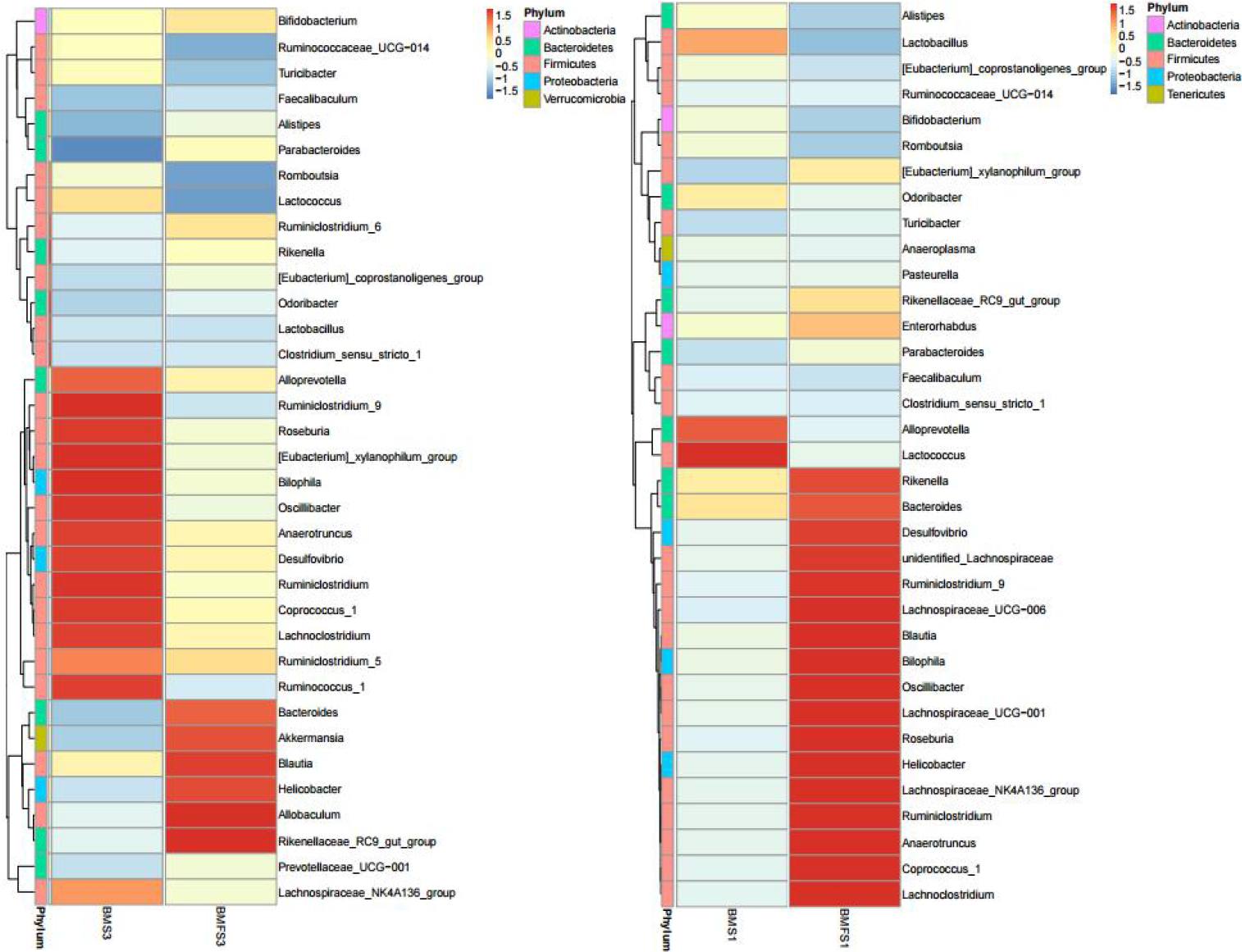
The ranked heat-maps of Top 35 bacterial genera in pre-modeling mice (a), modeling mice (b) and co-modeling MW drinking mice (c). BMS1: a fecal sample from a mouse drinking MW; BMFS1: a fecal sample from a mouse drinking BMW.

In such a context, BMW should not contain those functional competitors because of being boiled. Considering this situation, we determined to exclude CS-BC+BMW mice thereafter for further evaluation.

As listed in Table 6, almost all compared SSB/MEB and SRB, including *Bacteroides, Akkermansia*, *Bifidobacterium,* and *Desulfovibrio* are more abundant in CS-BC samples compared to AL samples. In contrast, almost all listed SSB/MEB and SRB are rare in CS-BC+MW samples. After MW drinking, *Akkermansia* even disappear, while *Bifidobacterium* are enriched in CS-BC+MW samples. This latter result implied that *Akkermansia* might be antagonized by *Bifidobacterium*.

**Table 6.** The proportion of SSB/MEB and SRB in the fecal samples from AL, CS, CS-BC and CS-BC+MW mice.

| SSB/SRB | Group | Model | Treatment |
| --- | --- | --- | --- |
| *Bacteroides* (containing SSB) | AL | 0.009 | 0.003 |
|  | CS-BC | 0.06 | 0.022 |
|  | CS-BC | 0.02 (-MW) | 0.03 (+MW) |
| *Akkermansia* (MEB) | AL | 0.00001 | 0.00001 |
|  | CS-BC | 0.007 | 0.0004 |
|  | CS-BC | 0 (-MW) | 0 (+MW) |
| *Bifidobacterium* (MEB) | AL | 0.01 | 0.0009 |
|  | CS-BC | 0.01 | 0.009 |

|  | CS-BC | 0.06 (-MW) | 0.02 (+MW) |
| --- | --- | --- | --- |
| *Desulfovibrio* (SRB) | AL | 0.001 | 0.002 |
|  | CS-BC | 0.006 | 0.02 |
|  | CS-BC | 0.001 (-MW) | 0.001 (+MW) |

Note: AL: *Ad libitum* chew-fed mice; CS: chondroitin sulfate-fed mice; CS-BC: chondroitin sulfate and *Bacillus cereus*-fed mice; CS-BC+MW: CS-BC mice treated by MW.

Interestingly, CS-BC mice also show decrease in the abundance of SSB/MEB and SRB, suggesting CS-BC feeding may nourish *Clostridia* (see above) and other bacteria, such as SSB/MEB and SRB. However, overgrowth of *Clostridia* may antagonize SSB/MEB and SRB because *Clostridia* are short-chain fatty acid (SCFA) producers, which may be not beneficial to the propagation of other bacteria.

We did not distinguish, from the data listed in Table 6, which one, BC or MW, acts as a chief effector in modulating SSB/MEB and SRB, but we believed that both BC and MW are most likely functioning because BC has been used as a prebiotic and MW can shape the gut microbiome. To avoid such an uncertainty, however, we added LPS mice for subsequent comparison because LPS injection would not be affected by gut bacteria.

### Suppression of LPS-triggered inflammation by co-modeling MW drinking

To monitor whether co-modeling MW drinking would affect the extent of endotoxinemia derived from LPS injection or CS-BC feeding, we determined the serum LPS levels in AL, LPS, CS-BC, and CS-BC+MW mice. Consequently, a higher serum level of LPS was determined in LPS mice than that of CS-BC+MW mice (Fig. 6a).

**Figure 6.**
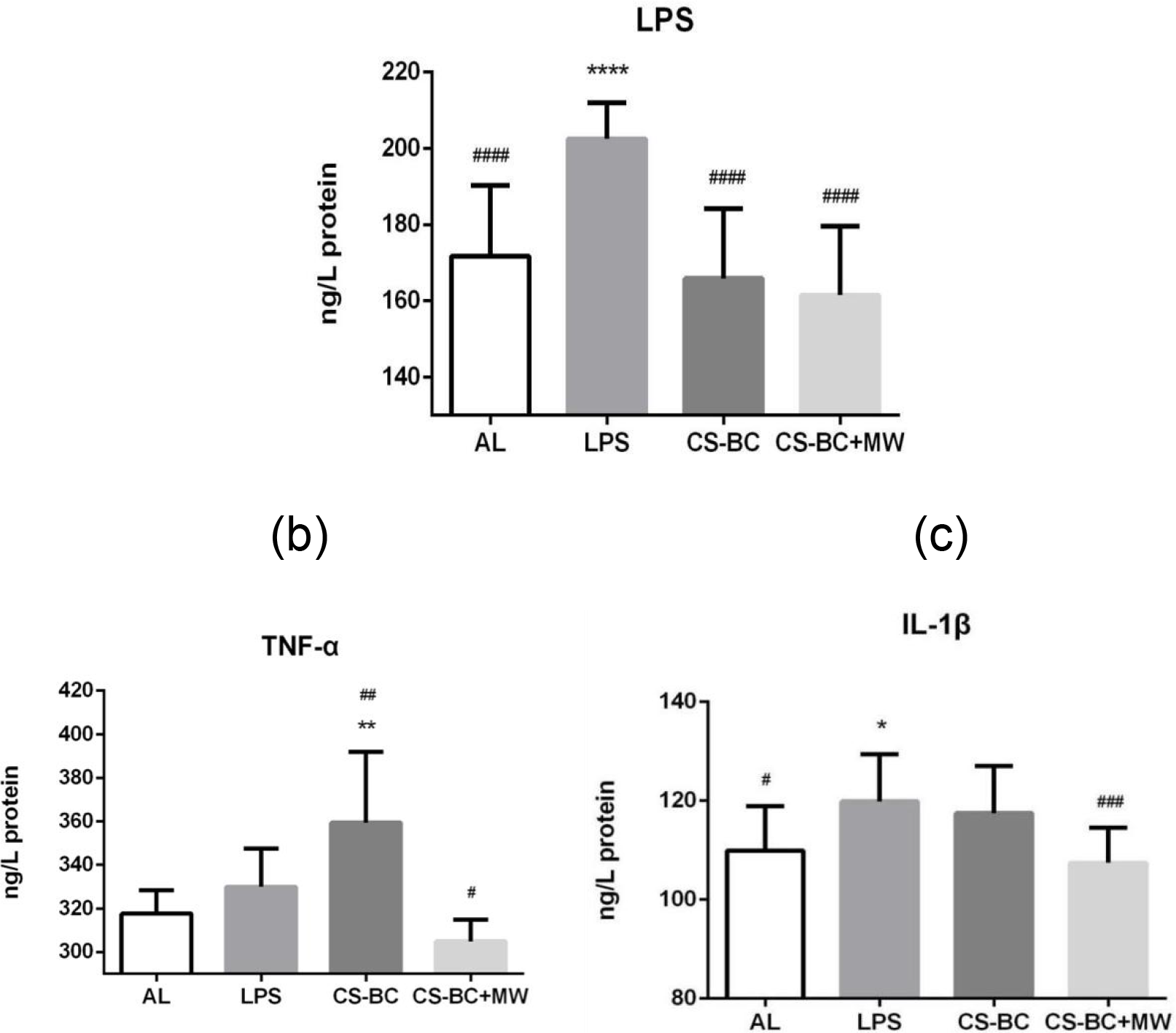
The levels of LPS and pro-inflammatory cytokines in AL, LPS, CS-BC, and CS-BC+MW mice. (a) The serum LPS levels (ng/L protein, *n*=8); (b) The serum TNF-α levels (ng/L protein, *n*=8); (c) The serum IL-1β levels (ng/L protein, *n*=8). AL: *Ad libitum* chew-fed mice; LPS: lipopolysaccharide-injected mice; CS-BC: chondroitin sulfate and *Bacillus cereus*-fed mice; CS-BC+MW: CS-BC mice treated by MW.

Accordingly, the levels of two major pro-inflammatory cytokines, TNF-α and IL-1β, are higher in LPS mice than CS-BC+MW mice (Fig. 6b and 6c), addressing that LPS triggers inflammation, whereas co-modeling MW drinking exerts nanti-inflammatory effects.

### Compromise of oxidative and hypoxic responses by co-modeling MW drinking

To further evaluate whether co-modeling MW drinking would modulate the inflammation-mediated oxidative stresses, we quantified the responsible mRNAs and encoded proteins for anti-oxidative responses. In consequences, *SOD2* mRNA that encodes the mitochondrial Mn-SOD that scavenges superoxide emitted from respiratory chains was found to exhibit a higher level in LPS mice than CS-BC+MW mice (Fig.7a-7b).

**Figure 7.**
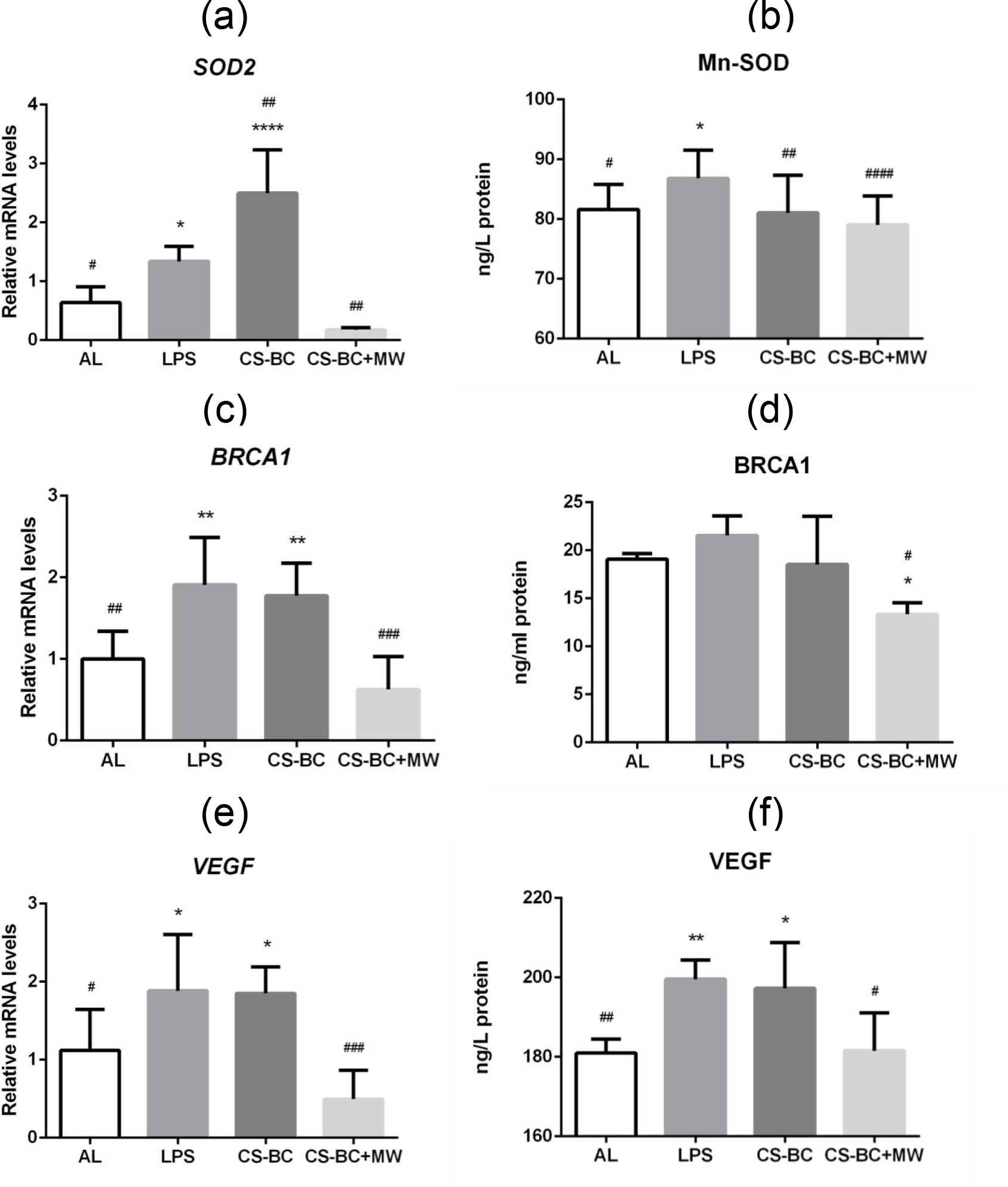
The expression levels of LPS-triggered oxidative and hypoxic response genes in AL, LPS, CS-BC, and CS-BC+MW mice. (a) The mammary *SOD2* mRNA levels (*n*=8); The mammary SOD2 levels (ng/L protein, *n*=8); (c) The mammary *BRCA1* mRNA levels (*n*=8); (d) The mammary BRCA1 levels (ng/L protein, *n*=8). (e) The mammary *VEGF* mRNA levels (*n*=8); (f) The mammary VEGF levels (ng/L protein, *n*=8). AL: *Ad libitum* chew-fed mice; LPS: lipopolysaccharide-injected mice; CS-BC: chondroitin sulfate and *Bacillus cereus*-fed mice; CS-BC+MW: CS-BC mice treated by MW.

Because of an attenuation of oxidative stresses mainly derived from the dysfunctional mitochondria, co-modeling MW drinking should also mitigate chromosomal DNA damage, thereby downregulating the DNA repairing responsible tumor suppressor *BRCA1* gene. Indeed, *BRCA1* mRNA and BRCA1 are synchronously declined in CS-BC+MW mice as compared with LPS mice (Fig. 7c-7d).

To evaluate the effects of co-modeling MW drinking on the inflammation-evoked hypoxic responses, we determined the levels of *VEGF* mRNA that encodes the angiogenesis-promoting factor VEGF. As expectation, both *VEGF* mRNA and VEGF levels are declined in CS-BC+MW mice as compared with LPS mice (Fig. 7e and 7f).

These results demonstrated that co-modeling MW drinking is sufficient to repress LPS-induced oxidative stresses, during which the negative impact of inflammation on the mitochondrial and chromosomal structure and function is abrogated upon *SOD2*/Mn-SOD- and BRCA1-mediated anti-oxidative and anti-mutagenic effects. While downregulation of *SOD2*/Mn-SOD represents a less extent of mitochondrial dysfunction, downregulation of *BRCA1* indicates a mild degree of chromosomal DNA damage.

### Downregulation of breast cancer markers and interruption of mammary hyperplasia by co-modeling MW drinking

To figure out how can co-modeling MW drinking affect the tumorigenesis, we determined the expression levels of breast cancer markers and examined the pathological changes of mammary glands in mice. As anticipation, the breast cancer cell proliferation-related *Ki67* mRNA and the TNBC-specific transcription factor-encoding *BCL11A* mRNA are lower in CS-BC+MW mice than those in LPS mice (Fig. 8a and 8b). Accordingly, mammary tubular hyperplasia occurs in LPS mice, but not appears in CS-BC+MW mice (Fig. 8c-8f), suggesting a positive correlation of mammary tubular hyperplasia with endotoxinemia, inflammation and hypoxia.

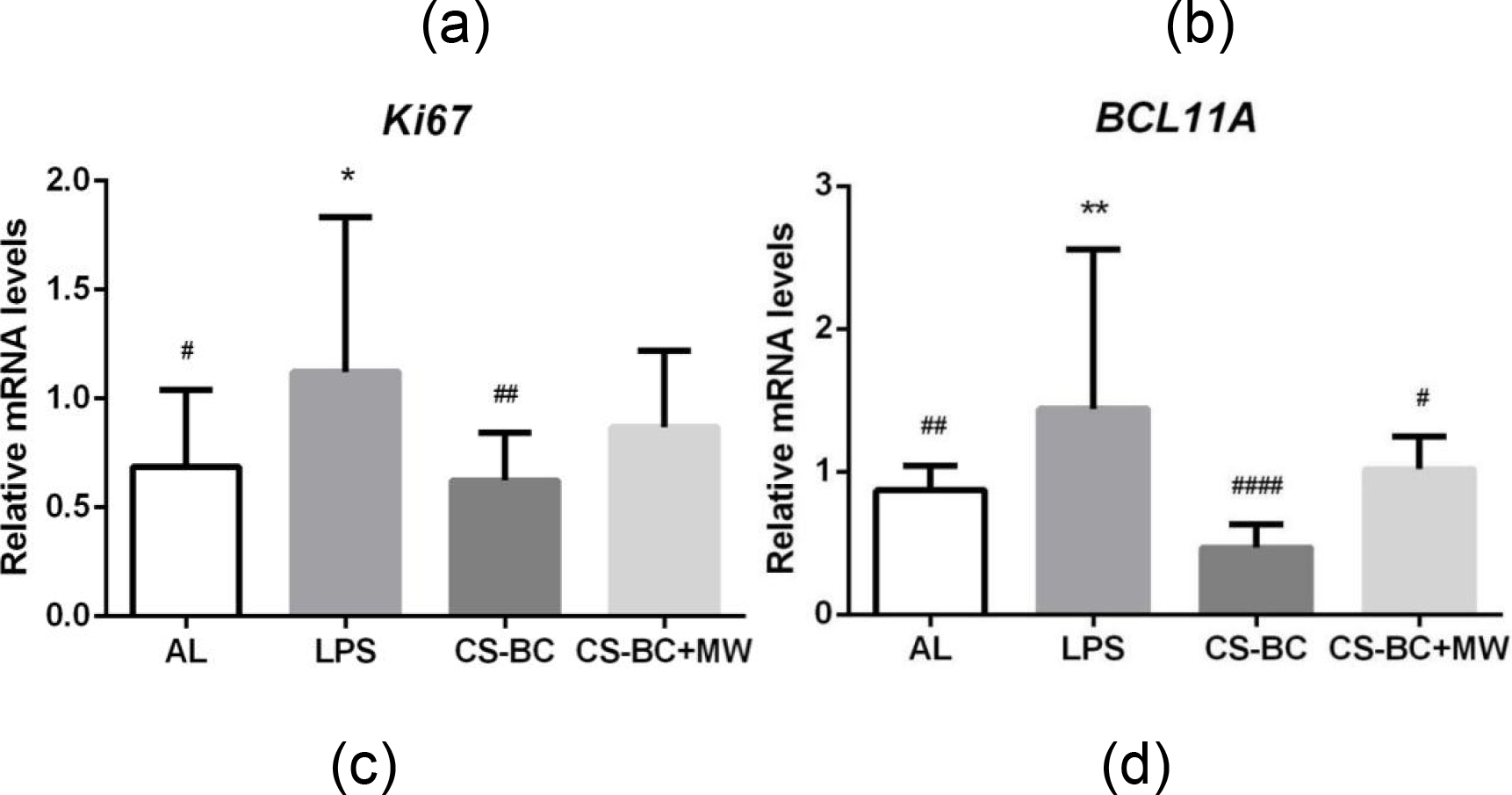

**Figure 8.**
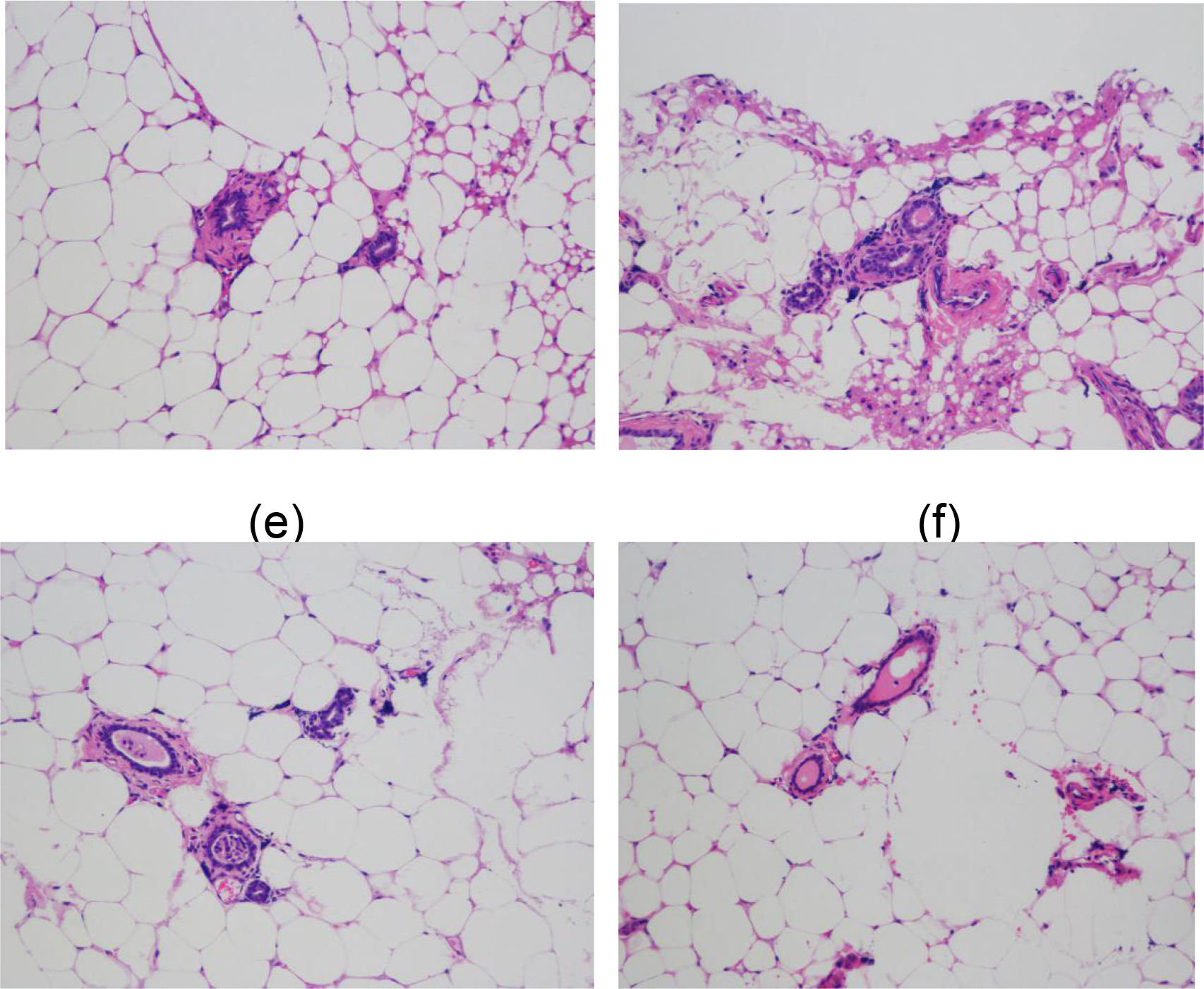
The quantification of breast cancer markers and the histochemical presentation of mammary hyperplasia in AL, LPS, CS-BC, and CS-BC+MW mice. (a) The mammary *Ki67* mRNA level; (b) The mammary *BCL11A* mRNA level; (c) The AL mammary tissue; (d) The LPS mammary tissue; (e) The CS-BC mammary tissue; (f) The CS-BC+MW mammary tissue. AL: *Ad libitum* chew-fed mice; LPS: lipopolysaccharide-injected mice; CS-BC: chondroitin sulfate and *Bacillus cereus*-fed mice; CS-BC+MW: CS-BC mice treated by MW.

These results demonstrated that co-modeling MW drinking is apt to repress the onset of early-phase breast cancer-like mammary hyperplasia by curing gut dysbiosis.

## Discussion

In Bama, 881 individuals who lived above 90 yeas old were registered among 269,800 residents in 2010 [20]. It was documented that longevity of Bama centenarians is correlated with the genetic difference, mainly the polymorphism of several related genes, such as apolipoprotein C-I [21], paraoxonase 1 [22], catechism-O-methyl transferase [23], methylene tetrahydrofolate reductase [24], peroxisome proliferator-activated receptor delta [25], cholesterol ester transfer protein [26], microsomal triglyceride transfer protein [27], and mitochondrial DNA [28]. Alternatively, dietary fibers [29] and mineral elements [30] were also attributed to longevity although the mechanisms involved are awaiting revelation.

The typical gut microbiota communities that are affected by the fiber-rich diets may link to lifespan extension in Bama residents. The age-related OTUs were characterized as *Ruminococcacea*, *Clostridiaceae*, and *Lachnospiraceae*, among which the former two are increased in Bama centenarians [31]. In regard to *Lactobacillus*, it was revealed that *L. salivarius* and *Weissella confusa* are increased, but *L. mucosae* are significantly decreased in the aged Bama citizens [32]. In the present study, we found that Bama elderly over 80 years even possesses the child-type gut microbiota profiles, characterized by more *Prevotella* and less *Bacteroides*. In contrast, travelers have an extraordinarily abundant *Bacteroides*, but no *Prevotella*.

A survey of the gut bacterial composition carried out in Burkina Faso of Africa indicated that *Prevotella* make up 53% of the gut bacteria in the rural children, but are totally absent in the gut of age-matched urban European children [33]. Another studies also demonstrated that long-term dietary patterns are strongly associated with the gut microbiome subtypes. For example, those who consume more protein and animal fats have dominant *Bacteroides*, while those who consume more carbohydrates, especially fibre, have dominant *Prevotella* [34].

It was observed that a shift of gut microbial floras toward a predominant *Bacteroides* community in older individuals compared to younger individuals [35]. As noted from a study, the microorganisms that are affected the most by aging are the diversity-associated taxa, comprising of *Prevotella* and associated genera, and their abundance is declined rapidly once individuals enter long-term care [36].

An association of MW with longevity, however, has not been concerned and investigated until currently. Even though it was proven that Nagoya hot MW (Takeda, Oita, Japan) can enrich the lean-related bacterial family, *Christensenellaceae* [17], no correlation of hot MW with lifespan has been explored up to now. By focused on the waterborne microbes, we disclosed for the first time that SSB/MEB and SRB are decreased in CS-BC+MW mice, but increased in CS-BC mice. This result strongly suggests that a disordered gut ecological system can be normalized by daily MW drinking.

For the prebiotic *Bifidobacterium*, its benefits to human health are commonly accepted [37]. Our results also indicated that either BC feeding or MW drinking increases the abundance of *Bifidobacterium*. On the other hand, *Akkermansia municinphila* is also considered to be beneficial to colonic health [38], yet its overgrowth was thought to be harmful because of inducing colitis [39]. We found that both BC and MW can deplete *Akkermansia*, suggesting that *Bifidobacterium* and *Akkermansia* may be antagonized each other.

We also reported that co-modeling MW drinking suppresses the overgrowth of *Clostridia* that are induced by CS-BC feeding. *Clostridia* are classified as sulfide-reducing bacteria that can be enriched accompanying with the harmful SSB and MEB. *C. difficile* is present in 2–5% of the human adult colon, but it may be opportunistically dominated when the normal balance of the gut flora homeostasis is disrupted by antibiotic therapies [40]. The risk of malnutrition is associated with increased microbiota diversity, particularly with a co-abundant *Clostradiales* subpopulation, and this subpopulation is also significantly associated with increased frailty [41]. Therefore, it might be said that the natural MW can confer health benefits to human, in part, by deprival of *Clostridia* in our gut.

## Conclusions

How to cope with opportunistic infection from the gut commensal bacteria remains unsolved in the elderly. In general, it would be realized by fecal microbiota transplantation [42], but more practical by probiotic supplementation [43] and dietary fiber consumption [44]. This study, showing a promise outcome of the natural and clean MW for the better control of gut bacteria, provides a convenient and safe choice to prevent the early onset of breast cancer by curating a high-fat diet- and red meat-induced gut dysbiosis [45, 46].

Interestingly, the female centenarians are more proportionally than the male centenarian in Bama, reflecting an effect of Bama MW-inhibited breast cancer on the lower mortality and longevity among the females. Our next work would be to evaluate the implications of post-modeling MW drinking as well as the heated but unboiled MW in the progression of breast cancer-like mammary hyperplasia in CS modeling mice.

The “Fountain of Youth”, Bama spring MW fulling with the non-pathogenic live bacteria, might have be found from the present study. Nevertheless, because bacteria-borne MW resources ubiquitously exist everywhere in the World, our results revealing that Bama MW compromises chromosomal damage and mitochondrial dysfunction would shed light into the eventual revelation of the secret of longevity in Bama as well as in the so-called “Blue Zone” districts [47].

## Declarations

### Acknowledgments

The authors thank Miss Ai-Fen You in TinyGene, Shanghai, China and Miss Fan Pei in Novogene, Beijing, China for assistance in the service of fecal sampling and bacterial sequencing.

### Availability of data and materials

All sequence count data can be found in Additional file 1-6.

### Authors’ contributions

JPS and QPZ conceived and designed the experiment. QX provided materials and discussed the experimental design. YPC, LLT, and DMC conducted animal experiments. YPC drawn column diagrams and performed statistical analysis. QPZ wrote the manuscript with input from all co-authors. All authors read and approved the final manuscript.

### Ethics approval and consent to participatsse

The animal experiments were approved by The Animal Care Welfare Committee of Guangzhou University of Chinese Medicine in China (No. SPF-2015009). The experimental protocols complies with the requirements of animal ethics issued in the Guide for the Care and Use of Laboratory Animals of the National Institute of Health (NIH) in USA.

